# Integrated analysis of sex-biased mRNA and miRNA expression profiles in the gonad of the discus fish *(Symphysodon aequifasciatus)*

**DOI:** 10.1101/492264

**Authors:** Yuanshuai Fu, Zhe Xu, Zaizhong Chen, Bin Wen, Jianzhong Gao

**Affiliations:** Key Laboratory of Freshwater Aquatic Genetic Resources, Ministry of Agriculture, Shanghai Ocean University, Shanghai 201306, China; Key Laboratory of Exploration and Utilization of Aquatic Genetic Resources, Ministry of Education, Shanghai Ocean University, Shanghai 201306, China; Shanghai Collaborative Innovation for Aquatic Animal Genetics and Breeding, Shanghai Ocean University, Shanghai 201306, China

**Keywords:** *Symphysodon aequifasciatus*, Gonad, Sex, mRNA, miRNA

## Abstract

The discus fish *(Symphysodon aequifasciatus)* is an ornamental fish that is well-known around the world. Phenotype investigation indicated that there are no significant differences in appearance between males and females of the discus fish. To better understand the sexual development mechanisms and obtain a high efficiency sex identification method in the artificial reproduction process of the discus fish, we constructed six cDNA libraries from three adult testes and three adult ovaries, and perform RNA-sequencing for identifying sex-biased candidate genes, microRNA (miRNA), and metabolic pathway using the Illumina Hiseq 4000. A total of 50,082 non-redundant genes (unigenes) were identified, of which 18,570 unigenes were significantly overexpressed in testes, and 11,182 unigenes were significantly overexpressed in ovaries, and 8 differentially expressed unigenes were validated by quantitative Real-Time PCR (qPCR). A total of 551 miRNAs were identified, of which 47 miRNAs were differentially expressed between testes and ovaries, and 7 differentially expressed miRNAs and one non-differential miRNA were validated by qPCR. Twenty-four of these differentially expressed miRNAs and their 15 predicted target genes constituted 41 important miRNA-mRNA interaction pairs, which may be important candidates for sex-related miRNAs and sex-related genes in the discus fish. Some of vital sex-related metabolic pathways were also identified that may play key roles in regulating gonad development of the discus fish. These results can provide important insights to better understand molecular mechanisms for sexual dimorphism in gonads development.

## Introduction

The discus fish *(S. aequifasciatus)*, one of the most demanded freshwater ornamental fish species around the world, has been widely cultured because of its brilliant colours and pretty disc-shaped body. With the improvement of people’s living standard, the demand for the discus fish is increasing. Before artificial reproduction, the discus fish must be matched by one male and one female. However, the male and female of discus fish cannot be judged according to the external features, which causes great difficulties in the mating between males and females in the artificial reproduction process. As a result, the rapid reproduction of the discus fish is supressed, its production is limited. Therefore, it is of great importance to establish artificial sex identification techniques that can be used in quick pairing of one male and one female for producing more and more progeny. A comprehensive understanding of developmental mechanism of the sexual dimorphism of *S. aequifasciatus* is urgently needed, including knowledge of the genes and miRNAs involved in the gonads of both sexes.

The factors underlying the decision to develop as one sex over the other are triggered by diverse cues among diverse species, such as sex chromosomes, the synergetic effects of multiple sex-associated genes, temperature, and social dynamics [1]. In fish, there is a higher variety of sex determination mechanisms, including genetic sex determination, environmental sex determination, or even interactions of genetic and environmental sex determination [2–5]. Gonad is an indispensable reproductive organ including testis and ovary, their development is controlled by many genes which are differentially expressed between testes and ovaries. To date, several master sex-determining genes have been identified and play a key role in regulating sex development as transcription factors. *SRY* in mammals [6] and *DMY/dmrt1bY* in medaka [7, 8] both initiate male sex determination, are required for testis formation in XY embryos and are sufficient to induce testis differentiation in XX embryos [9]. In females, there exists a number of essential ovary-specific genes, for example, *β-catenin, follistatin, FOXL2, R-spondin* and *WNT4* [9]. Deficiency of these genes may cause ovarian development stasis, and mutation of these genes may result in aberrant ovary development. Gametogenesis can be differentiated in spermatogenesis and oogenesis [10]. There are numerous gene expressions associated with gametogenesis in the reproductive stage in the mature gonads of teleosts [11, 12]. Previous studies have indicated that several key genes associated with steroid hormone enzymes are expressed during the stage of gametogenesis in tilapia [13]. We assumed that some of genes play key roles in regulating gonads development. Targeted differential expression analysis of genes and miRNAs between testes and ovaries will help us identify the molecular basis of testis development and ovary development in *S. aequifasciatus*.

MicroRNAs (miRNAs) are endogenously expressed, small non-coding RNAs that are approximately 18–22 nucleotides (nt) long and that post-transcriptionally regulate gene expression, inhibit targeted gene translation or degrade target mRNA by partial or complete complementarity binding to the 3’ untranslated region (UTR) of target genes in both animals and plants [14, 15]. The total set of transcripts (mRNA and non-coding RNA) involved in the transcriptome are transcribed at a specific organization during a particular developmental stage [16]. In organism, one miRNA may control the expression of several genes, or the expression of a single gene requires multiple miRNAs to work simultaneously [17]. Previous studies showed that miRNA may be an inducible factor to increase the complexity of organism with their roles in regulating gene expression [18–20]. In tilapia gonads, 111 differentially expressed miRNAs were identified between testes and ovaries, and the targets of these sex-biased miRNAs contained key genes encoding enzymes in steroid hormone biosynthesis pathways [21]. In rainbow trout, a total of 13 differential expression miRNAs were observed during stages of oogenesis, which indicated that they might regulate female gamete formation during oogenesis [22]. There is no available information on miRNAs in the testes and ovaries of the discus fish.

In this study, we are aiming to screen differentially expressed genes and miRNAs between testes and ovaries through RNA-sequencing, and identify key genes and miRNAs capable of regulating gonadal development or sex determination. This work will help to further understand the underlying molecular mechanisms of sex differentiation and sex determination in *S. aequifasciatus*.

## Materials and Methods

### Ethical statement

All fish used in this study were housed and sacrificed according to approved animal use protocols dictated by the Institutional Animal Care and Use Committee at the Shanghai Ocean University.

### Sample collection and RNA extraction

Three one-year-old males (255 ± 8.64 g) and three one-year-old females (228± 4.76 g), were obtained from the Ornamental Aquatic Breeding Laboratory in the College of Fisheries and Life Science of Shanghai Ocean University (Shanghai, China). The system (equipped with a volume of 120 L) temperature was maintained at 28 ± 0.5°C. The fish were fed with beef heart. The experimental fish was anaesthetized in well-aerated water containing a 100 mg/L concentration of Tricaine methanesulphonate (MS-222) (Argent Chemical Laboratories, Redmond, WA, USA). Ovary and testis samples were collected and immersed into liquid nitrogen and then stored at −80 °C until subsequent RNA isolation.

### RNA isolation

Total RNA was extracted using the miRNeasy Kit (QIAGEN, USA) according to the manufacturer’s protocol, and treated with RNase-free DNase I (TianGen, Beijing, China) to remove genomic DNA contamination. The total RNA quantity and purity were analysed by Bioanalyzer 2100 (Agilent, Technologies, Inc.) with RNA integrity number (RIN) numbers > 7.0. The concentration and quality of the purified RNA samples were determined utilizing a Nanodrop 2000C spectrophotometer (Thermo Scientific, Waltham, MA, USA) and RNA integrity was detected by agarose-gel electrophoresis, and A260/A280 ratios were between 2.0 and 1.9.

### Library construction and sequencing for mRNA

Six gonadal samples for mRNA transcriptome analysis were prepared using a Truseq^TM^ RNA Sample Prep Kit (Illumina, San Diego, USA) according to the manufacturer instructions. These mRNAs were isolated from >5 μg of gonadal total RNAs using oligo (dT) magnetic beads. These short fragment RNAs were transcribed to create first-strand cDNAs using random hexamer-primers, the second-strand cDNAs were then synthesized using RNase H, buffer, dNTP, and DNA polymerase I. These double stranded cDNAs were purified using Takara’s PCR extraction Kit (Takara Bio, Inc.), and then ligated with sequencing adapters, and resolved by agarose gel electrophoresis. Proper fragments were selected and purified and subsequently amplified by 15 cycles of PCR to create the cDNA libraries. The DSN kit (Evrogen, Russia) was used to normalize the cDNA libraries. These normalized cDNA libraries were sequenced on an Illumina Hiseq2500 sequencing platform, with 125 nt reads length and both end sequencing pattern.

### Library construction and sequencing for small RNA

Six gonadal samples for miRNA transcriptome analysis were prepared using a TruSeq^TM^ Small RNA Sample Prep Kit (Illumina, San Diego, USA) according to the manufacturer instructions. Small RNA was isolated from gonadal total RNA, and was ligated with proprietary 5’ and 3’adapter. Adaptor-ligated small RNAs were then reverse transcribed to create cDNA constructs using Superscript reverse transcriptase (Invitrogen, CA, USA). These generated small cDNA libraries were amplified by 15 cycles of PCR using Illumina small RNA primer set and Phusion polymerase (New England Lab, USA), and purified on a 6% Novex TBE PAGE gel. The purified PCR libraries were sequenced on an Illumina Hiseq2500 sequencing platform, with 50 sequencing cycle number, 50 nt reads length and single end sequencing pattern.

### Bioinformatic analysis of mRNA and miRNA transcriptome data

The clean reads for mRNA transcriptome were obtained using NGS QC TOOLKIT v2.3.3 software [23] by filtering out adapter sequences, low quality reads (reads with ambiguous bases ‘N’), and reads with more than 10% Q < 25 bases. The clean reads were assembled into non-redundant transcripts using Trinity program (http://trinityrnaseq.sourceforge.net/) [24] with default K-mers = 25. The non-redundant transcripts less than 100bp in length and partially overlapping sequence were removed. Next, the remained non-redundant transcripts were annotated by Blast search against the NR protein, the GO, COG, and KEGG database using an E-value cut-off of 10^-5^. The functional annotation by GO terms [25] was carried out using Blast2go software, the functional annotation against the COG [26] and KEGG database [27] was performed using Blast software.

The raw reads from miRNA transcriptome were subjected to initially filter and remove low quality reads (including reads shorter than 18 nucleotide) and adapter sequences. The clean reads with a length of 18~ 26 nucleotide were subsequently aligned to Rfam 11.0 and the NCBI database searching, and these reads with similar to rRNA, tRNA, snRNA, snoRNA, and scRNA were removed. The remnant reads were aligned against correlation sequences in miRBase 21[28] and the reference genome of *S. aequifasciatus* non-redundant transcripts of this study, allowing length variation at both 3’ and 5’ ends and one mismatch inside of the sequence [29]. The unmapped sequences were BLASTed against the specific genomes, and the hairpin RNA structures containing sequences were predicated from the flank 80 nt sequences using RNAfold software (http://rna.tbi.univie.ac.at/cgi-bin/RNAfold.cgi).

Two computational target prediction algorithms (TargetScan and miRanda) were used to predict the target genes of miRNA from blast matching against the *S. aequifasciatus* mRNA transcriptome sequence of this study. miRanda was used to match the entire miRNA sequences. The TargetScan [30] parameters were set as a context score percentile > 50. The miRanda parameters [31] were set as free energy < −10 kcal/mol and a score > 50. All miRNA targets were categorized into functional classes using the GO terms and KEGG pathway. And the results predicted by the two algorithms were combined and the overlaps were calculated.

### Expression analysis of mRNAs and miRNAs

mRNA expression level between different mRNA transcriptome were measured by RPKM (reads per kb per million reads) using RSEM software [32, 33]. All transcript sequences obtained by splicing in Trinity were used as reference sequences. Each sample’s Valid Data alignment was quantified to the reference sequence. The main parameters were: no-mixed, no-discordant gbar 1000, and end-to-end-k200. The mRNAs with log-fold difference ratios (log2Ratio) ≥ 1 and false discovery rate (FDR< 0.05) were considered to be significantly differentially expressed.

miRNA differential expression based on normalized deep-sequencing counts was analysed by t-test. Comparisons between testes and ovaries were made to identify significantly differentially expressed miRNAs (|log2(fold change)|> 1 and P-value≤ 0.05). Data normalization followed the procedures as described in a previous study [34].

### Quantitative real time PCR validation for miRNA and mRNA expression

To validate the RNA-Seq data, 8 mRNAs and 8 miRNAs were randomly selected from differentially expressed mRNAs and miRNAs, and specific primers for mRNAs, miRNAs, 18S rRNA and 5S rRNA (Supplementary Table S1), were designed to quantify their expression levels between ovaries and testes using Quantitative real time PCR (qRT-PCR). Total RNAs were reverse transcribed to cDNAs using M-MLV Reverse Transcriptase (Promega), with an equal amount of mixed reverse primer of the Oligo(dT)_18_ (Takara) and random primer (hexadeoxyribonucleotide mixture; pd(N)6) (Takara) for the quantification of mRNAs, and stem-loop RT primer (Supplementary Table S1) for the quantification of miRNAs.

qRT-PCR was performed using PowerUp™ SYBR^TM^ Green Master Mix (Thermo Fisher Scientific Inc., IL, USA) on a CFX96 Touch™ Real-Time PCR Detection System (Bio-Rad, USA). The 18s rRNA that was not differentially expressed between testes and ovaries, was used as an internal control for mRNA expression levels, and the 5s rRNA without differential expression between testes and ovaries was used as the internal control for the normalization of miRNA expression levels. qRT-PCR reactions of 20 μl contained: 10.0μ1 SYBRTM Green Master Mix, 1.0μl forward primer and 1.0μl reverse primer, 1.0μl cDNA, and 7.0μl DEPC H_2_O, and the amplification procedure was carried out at 95°C for 2 min, 40 cycles of 95°C for 10 s and 60°C for 20 s followed by disassociation curve analysis. Samples were replicated two times for each run, and the average Ct value used to calculate gene expression levels. The relative expression levels were determined using the 2^-∆∆Ct^ method (Livak and Schmittgen, 2012). All primers were listed in Supplementary Table S1.

### Statistical analysis

All of data are expressed as the mean ± SEM. The level of significance was analyzed by one-way analysis of variance (ANOVA) with SigmaPlot 12.5 and SigmaPlot 3.5 software. Statistically significant differences were examined by paired t-test. A value of *p* < 0.05 was considered to be statistical significance.

## Results

### De novo assembly and functional annotation of mRNA transcriptome

Six cDNA libraries from three testes and three ovaries were sequenced using Illumina Hiseq 4000 sequencing. A total of 437,621,708 reads were obtained from six cDNA libraries (Table 1). After quality filtering using the Trinity de novo assembly method, we obtained 50,082 non-redundant genes (unigenes) with an average length of 885 bp and 65,496 transcripts with an average length of 1,077 bp (Table 2). All of the raw mRNA transcriptome sequencing data have been submitted to the NCBI database (https://www.ncbi.nlm.nih.gov/sra/SRP148426).

**Table 1.** Summary of Illumina Hiseq 4000 sequence reads.

| **Sample** | **raw data reads** | **Valid Data reads** | **Q20**<br>**percentage** | **Q30**<br>**percentage** | **GC**<br>**percentage** |
| --- | --- | --- | --- | --- | --- |
| Testis_1 | 88,682,568 | 86,322,782 | 97.26 | 92.56 | 50.25 |
| Testis_2 | 82,098,772 | 79,904,932 | 97.39 | 92.72 | 50.72 |
| Testis_3 | 54,797,130 | 53,361,960 | 97.28 | 92.51 | 49.13 |
| Ovary_1 | 69,045,830 | 66,995,128 | 96.93 | 91.72 | 50.51 |
| Ovary_2 | 62,969,370 | 61,204,582 | 97.27 | 92.41 | 51.22 |
| Ovary_3 | 80,028,038 | 77,880,556 | 97.38 | 92.70 | 49.97 |

**Table 2.** De novo assembly statistics of the discus fish transcriptomic sequences.

| **Category** | **All** | **Median GC%** | **Mean GC%** | **Mean Length** | **N50 of contigs** |
| --- | --- | --- | --- | --- | --- |
| gene | 50,082 | 47.80 | 47.28 | 885 | 1,540 |
| transcript | 65,496 | 48.10 | 47.64 | 1,077 | 1,896 |

To annotate and analyse the putative functional roles of unigenes, all unigenes were compared with the Swiss-prot database, Pfam database, and NCBI non-redundant nucleotide sequences (Nr) in the priority order of Gene ontology (GO), eukaryotic Orthologue Groups (KOG) database and Kyoto Encyclopaedia of Genes and Genomes (KEGG) database. A total of 25,026 (24.51%) unigenes were annotated in the NR database, and 22739 unigenes in the Swiss-prot, while the other unannotated unigenes represent novel genes of unknown functions (Table 3). A structured and controlled vocabulary to describe gene products were obtained using GO and KOG analysis. A total of 20,176 (40.29%) unigenes were assigned to the GO database (Table 3). Three distinct GO categories were characterized with 10 molecular function groups, 15 cellular component groups, and 25 biological process groups (Supplementary Figure 1). A total of 21,311 (42.55%) unigenes were assigned to the KOG database and classified into 25 functional categories (Supplementary Figure 2). KEGG analysis was performed to identify potential candidate unigenes in biological pathways. A total of 15,702 (31.35%) unigenes were assigned to the KEGG database (Table 3), and mapped onto 36 predicted pathways (Supplementary Figure 3).

**Table 3.** Annotation statistics of the discus fish transcriptome sequencing.

| **gene_number** | **Swiss-prot** | **Pfam** | **Nr** | **GO** | **KOG** | **KEGG** |
| --- | --- | --- | --- | --- | --- | --- |
| 50,082 | 22,739 | 20,450 | 23,371 | 20,176 | 21,311 | 15,702 |
| 100% | 45.40% | 40.83% | 46.67% | 40.29% | 42.55% | 31.35% |

### Differentially expressed mRNAs

A total of 29,752 differentially expressed genes (DEGs) were detected by comparing DEGs between testis and ovary transcriptomic sequencing (Supplementary Table S2). A total of 18,570 of DEGs were higher expressed and 11,182 lower expressed in the testis compared to the ovary (Fig. 1A). There were 3,151 genes that were specifically expressed in the testis, 222 in the ovary and 26,379 in both tissues. We can visually find the DEGs by a Volcano plot (Fig. 1B). The X-axis shows the differences in expression given as the log values, while the Y-axis shows the significant differences in expression as negative log values. DEGs are indicated by red dots and non-differentially expressed genes are indicated by blue dots.

**Fig 1.**
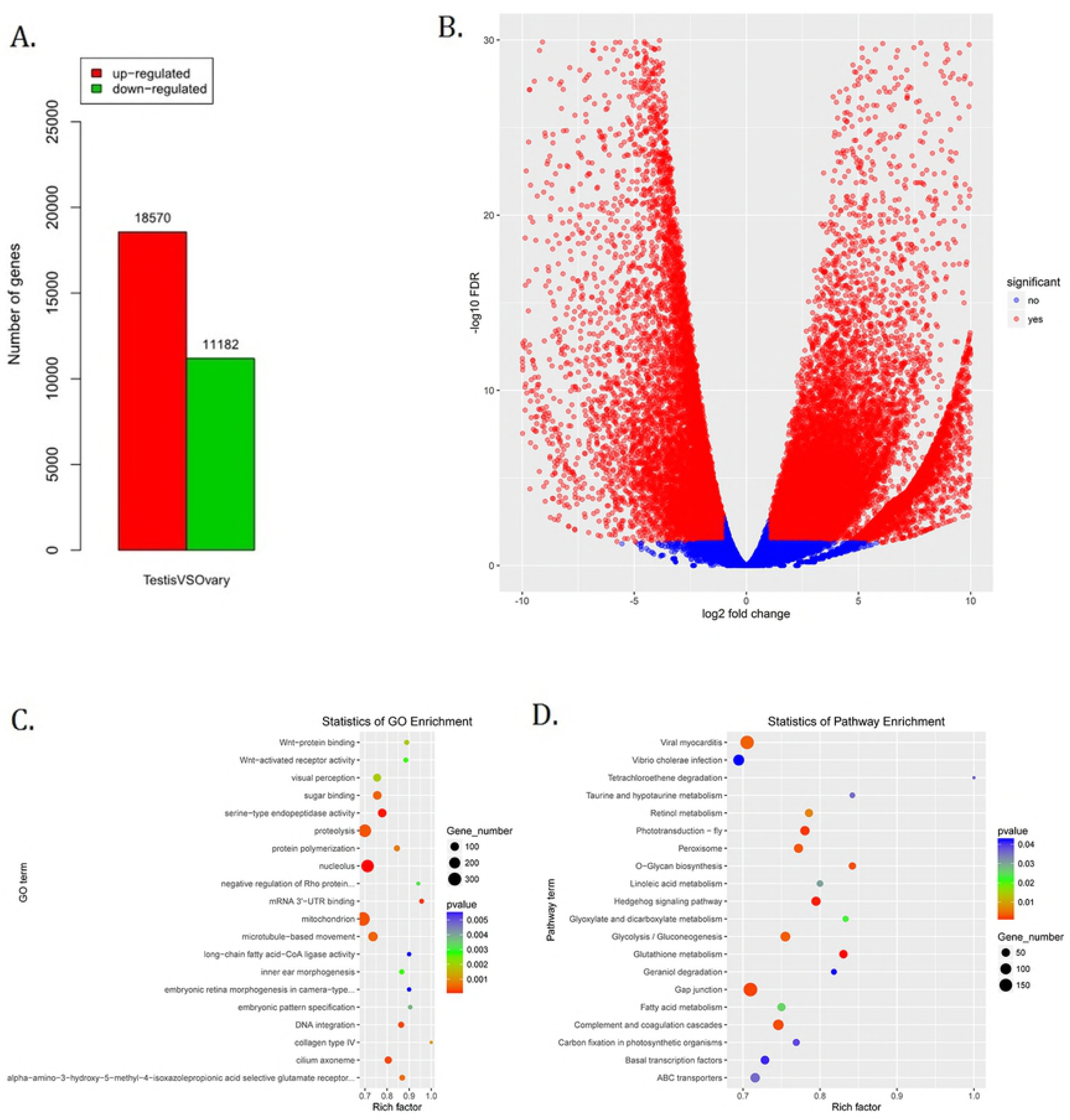
Identification, GO, and pathway of differentially expressed genes between testis and ovary. (A). Number of up/down expressed mRNAs in testis VS ovary. The ‘up-regulated’ means that these unigenes were higher expressed in the testes comparing to the ovaries, and the ‘down-regulated’ means that these unigenes were lower expressed in the testes comparing to the ovaries; (B). Volcano plot of different expression genes between the testis and ovary; (C). GO scatter diagram different expression genes between the testis and ovary; (D). KEGG scatter diagram different expression genes between the testis and ovary.

To further determine and compare the functions of these differentially expressed genes, a total of 122 GO terms with including these differentially expressed genes, were classified into 3 gene ontology (GO) categories (cellular component, biological process and molecular function) (Supplementary Table S3, Fig. 1. C). The DEGs were compared to the KEGG pathway to gain an overview of the gene pathway networks. A total of 1,251 DEGs involved in 243 pathways were predicted in the pairwise comparison of testis-vs-ovary *(p* < 0.05) (Supplementary Table S4, Fig. 1. D).

### Differentially expressed miRNAs

In this study, a total of 551 miRNAs were identified from the gonad tissues, and they ranged from 18–26 nt in length (Table 4). By comparing miRNA expression levels between testes and ovaries, a total of 47 miRNAs were differentially expressed between testis and ovary (Supplementary Table S5). 31 miRNAs were significantly higher expressed and 16 miRNAs significantly lower expressed in the testes comparing to ovaries (Fig. 2A). We found that four miRNAs were specifically expressed in the testis, and they were dre-miR-124-5p, ccr-miR-7133, PC-5p-60222, aca-miR-726, while three miRNAs were specifically expressed in the ovaries. Using the heat map of the differentially expressed miRNAs, one can intuitively see the changes in miRNA expression (Fig. 2B). These differentially expressed miRNAs were regarded as candidate sex-related miRNAs.

**Table 4.**
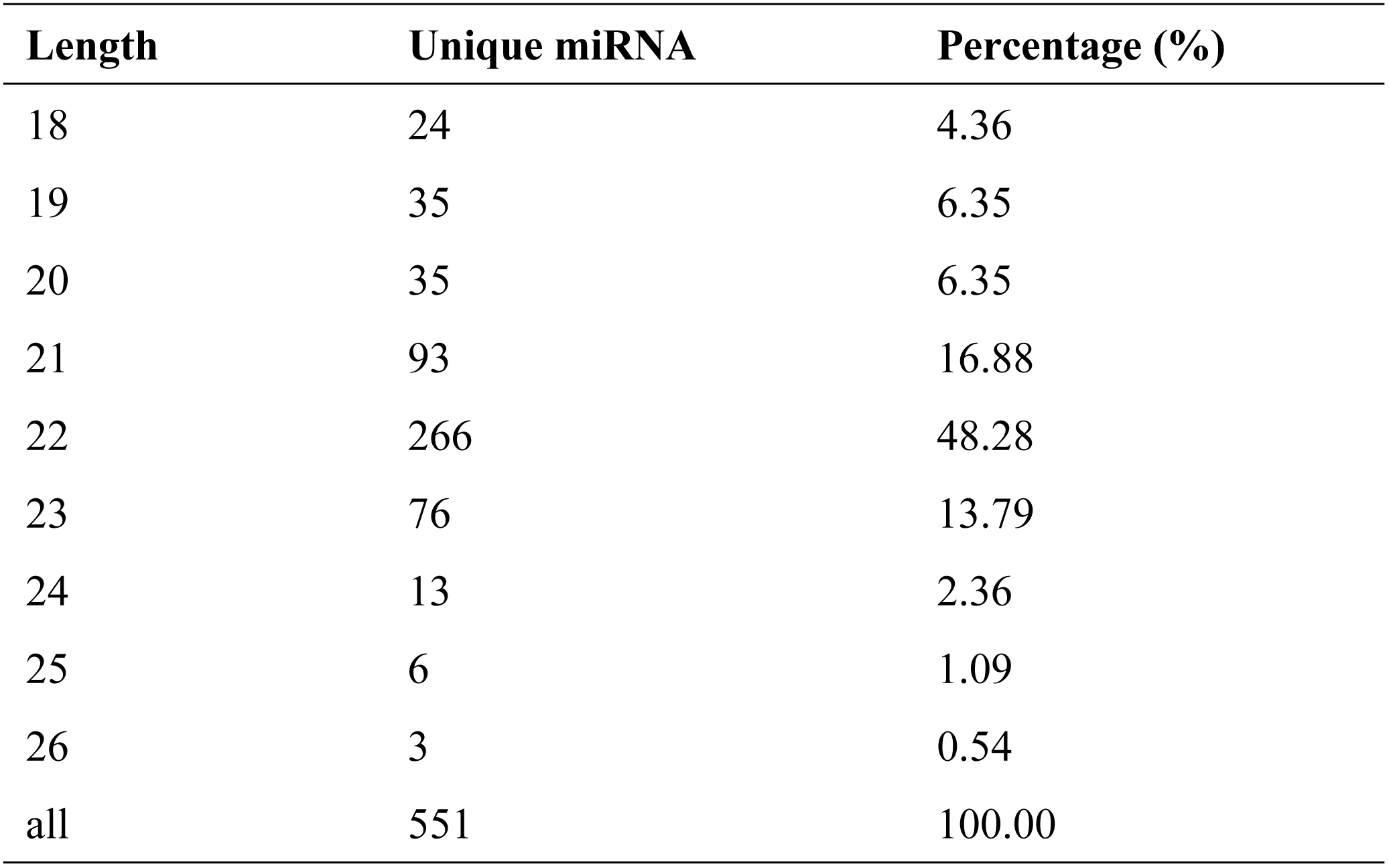
Distribution of these identified miRNAs in length.

**Fig 2.**
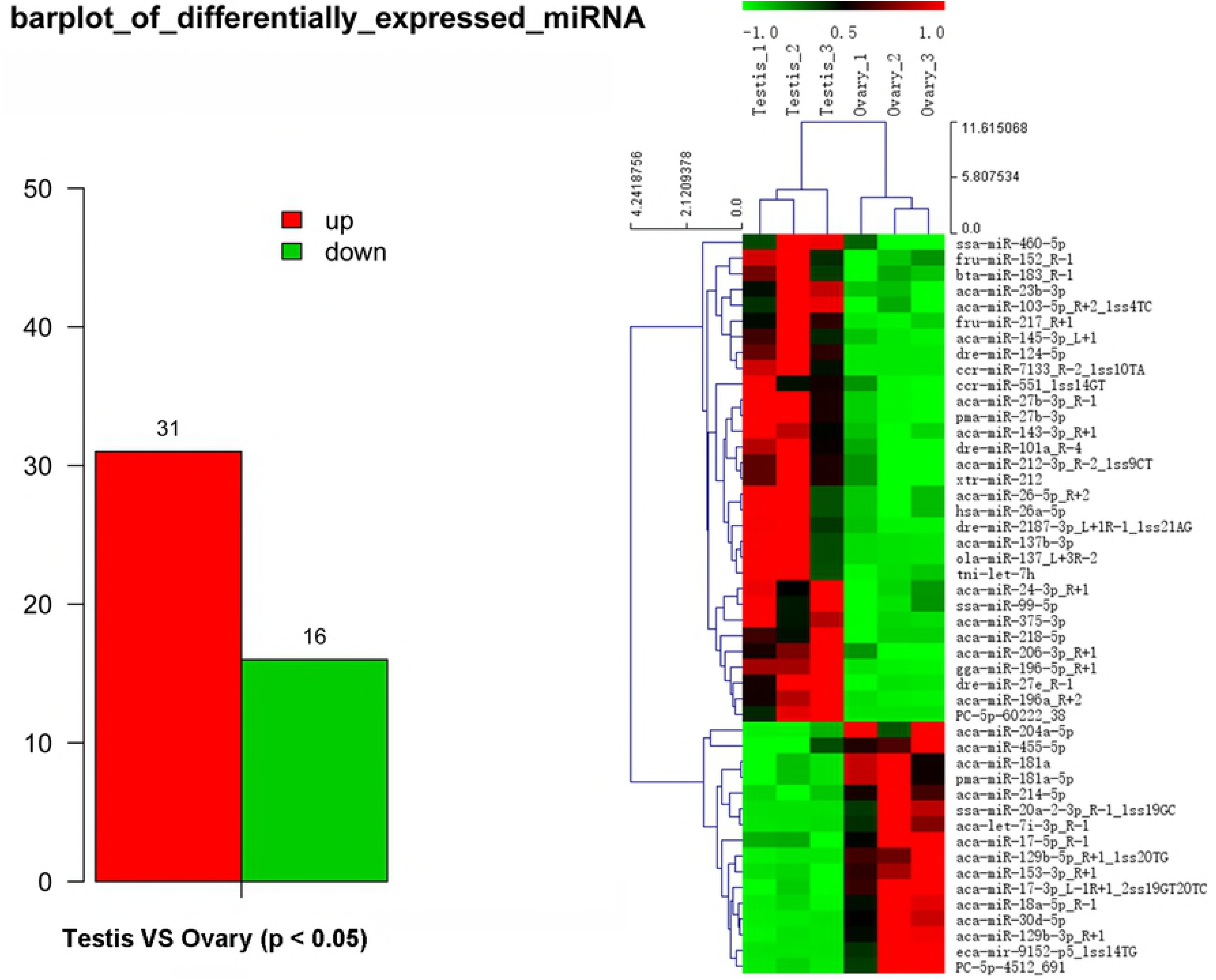
Differential expressed miRNAs between testis and ovary. (A). Number of up/down expressed miRNAs in testis and ovary. The ‘up’ means that these miRNAs were higher expressed in the testes comparing to the ovaries, and the ‘down’ means that these miRNAs were lower expressed in the testes comparing to the ovaries; (B). Heat map of the differentially expressed miRNAs in testis and ovary.

### Integrated analysis of DEMs and DEGs

A total of 9, 311 differentially expressed genes under the control of 47 miRNAs were identified, and of these, 5,600 were positively regulated and 3,711 were negatively regulated. In the KEGG functional enrichment, by percentage, we identified the KEGG pathway that needs to be focused on in the joint analysis (Fig. 3A). Approximately 41 DEGs were assigned to 37 signal pathways (Fig. 3B), and DEMs were involved in 23 signal pathways (Fig. 3C). We identified 41 miRNA-mRNA interaction pairs with 24 differentially expressed miRNAs targeting 15 DEGs. (Supplementary Table S7).

**Fig 3.**
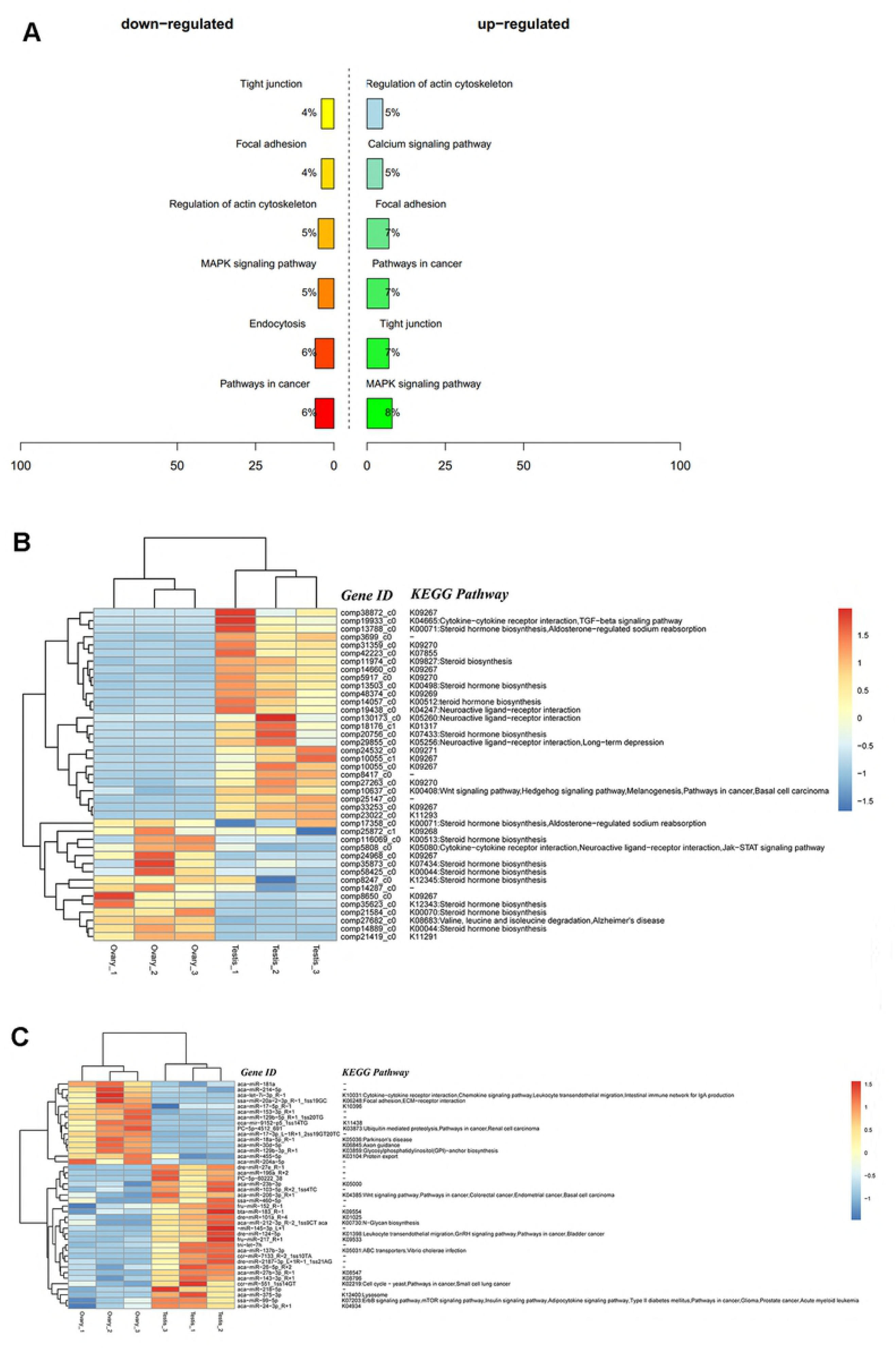
Integrated analysis results of DEMs and DEGs. (A) Precent of up/down expressed mRNAs associated with differentially expressed miRNAs in KEGG enrichment. The ‘up-regulated’ means that these unigenes were higher expressed in the testes comparing to the ovaries, and the ‘down-regulated’ means that these unigenes were lower expressed in the testes comparing to the ovaries; (B) Clustering and pathway distribution of 41 differentially expressed mRNAs; (C) Clustering and pathway distribution of 41 differentially expressed miRNAs.

Different expression of miRNAs with their predicted target genes were explored for cognate mRNA targets in their respective unigenes list in order to profile miRNA-mRNA functional interactions. These identified 41 miRNA-mRNA pairs were involved in sex differentiation and steroid hormone biosynthesis (Supplementary Table S7). Among these candidate miRNA-mRNA interaction pairs, each miRNA can target one or more genes, while each gene can be targeted by one or more miRNAs, which indicated that there were complex regulatory networks between miRNAs and gene mRNAs (Fig 4). For example, miR-2187-3p, miR-24-3p, miR-181a, miR-181-5p, miR-101a, miR-218-5p, miR-217 and miR-7133) can target *Hsd11β2*, while *Dmrt1* was targeted by miR-129-5p, miR-27b-3p, miR-27e, miR-152, miR-27b-3p, miR-17-3p and miR-460.

**Fig 4.**
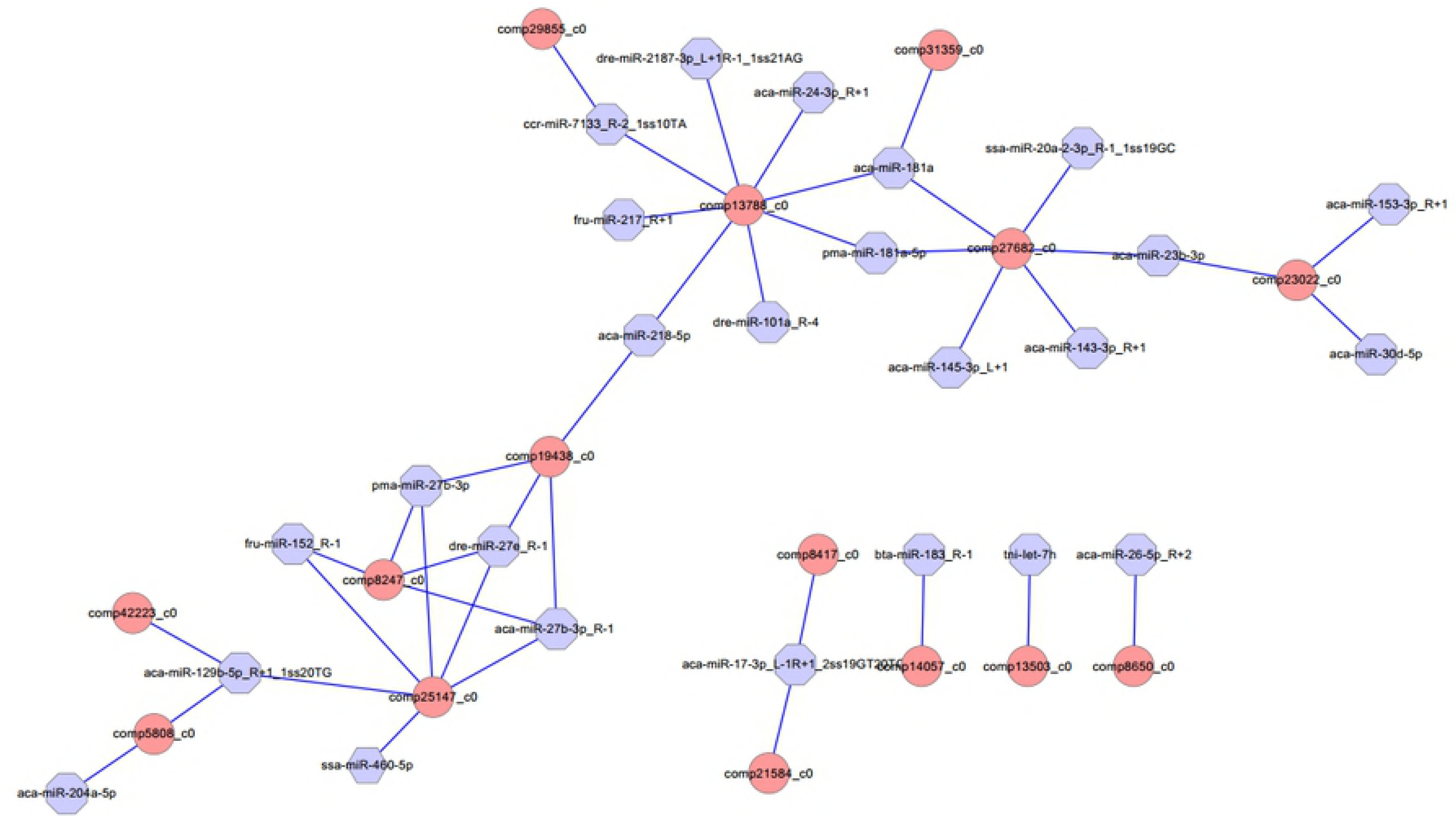
Proposed network of putative interactions between miRNAs and mRNAs in the gonad development. The regulation network of miRNAs and mRNAs involved in sexual development and maintenance is illustrated by Cytoscape. Pink represent miRNAs, and blue indicate their target genes.

### Validation of miRNA and mRNA expression using qRT-PCR

To verify the reliability and accuracy of the RNA-Seq results, we randomly selected 8 genes and 8 miRNAs to examine their expression levels between testis and ovary using qRT-PCR (Supplementary Table S6, Fig 5). Expression results of genes indicated that *Amh*, *Cyp11a1, Cyp17a1, Dmrt1*, *Dmrt2, Hira* and *Hsd11b2* were more highly expressed in the testis, while *Sox3* was more highly expressed in the ovary. Expression analysis of miRNAs indicated that miR-196a, miR-26a-5p and miR-375-3p were up-regulated, and miR-129b-5p, miR-18a-5p, miR-30d-5p and let-7i-3p were down-regulated. The qRT-PCR results for these 8 miRNAs and 8 mRNAs were similar to the RNA-Seq data.

**Fig 5.**
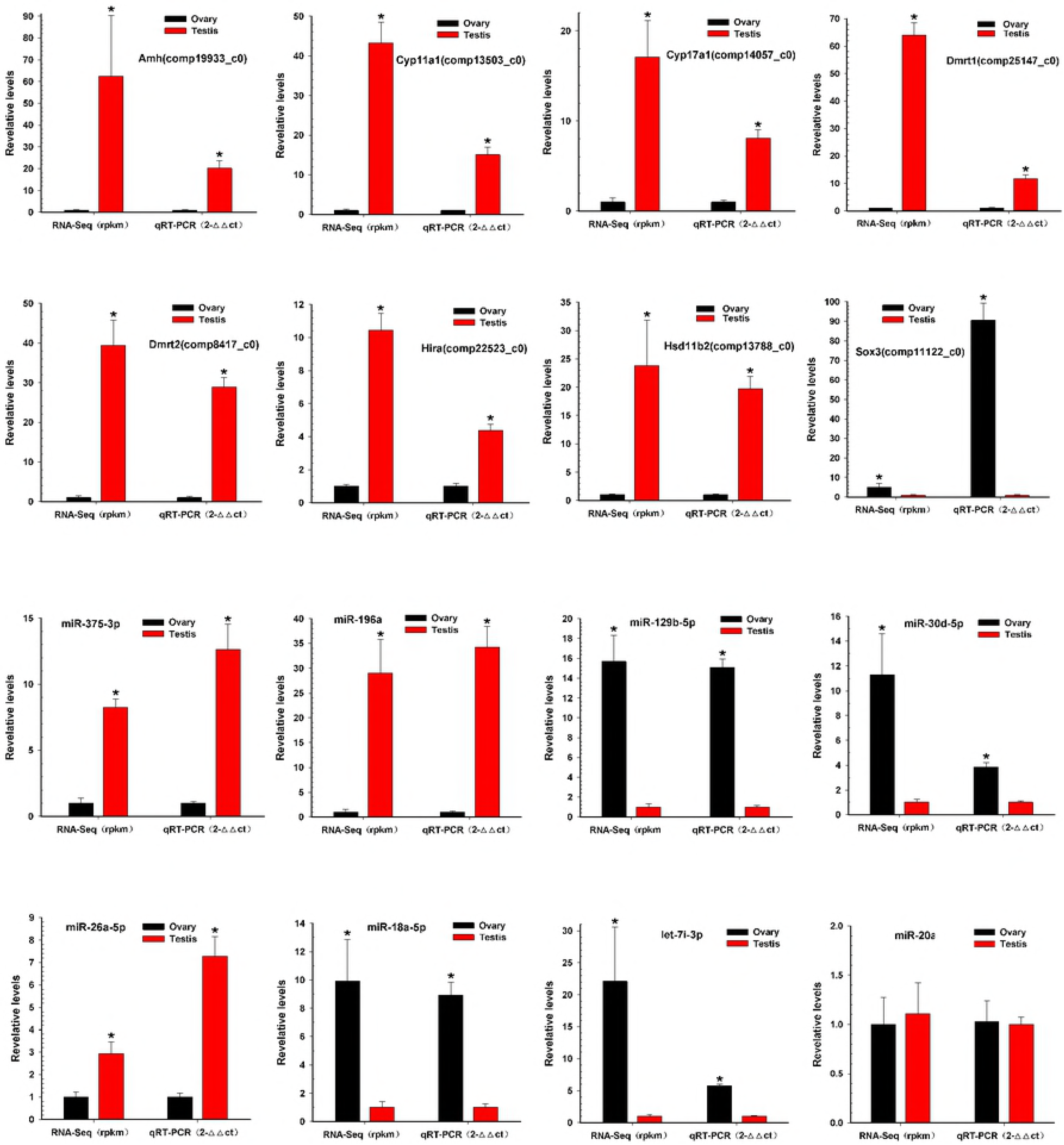
Validation of miRNAs and mRNAs expression between testis and ovary. QRT-PCR analysis of 8 gene mRNAs and 8 miRNAs selected randomly from RNA-Seq results between testis and ovary. All data were shown as mean ± SD. The *P < 0.05 mean a statistically significant difference.

## Discussion

To date, there is no available information regarding the mRNA transcriptome and miRNA in *S. aequifasciatus*. In this study, using a high-throughput sequencing approach, we performed RNA-Seq analysis to study the expression of miRNAs and mRNAs in the gonads. The aim of the present work was to identify differentially expressed mRNAs and miRNAs between testes and ovaries and to predict miRNA possible targets, as well as to discover possible mechanisms responsible for sexual dimorphism in gonads.

In this study, De novo assembly revealed 50,082 non-redundant genes in the gonad. Based on the functional annotation of non-redundant genes, several sex-related metabolic pathways were summarized from other species. The main metabolic pathways of steroid hormone biosynthesis and oocyte meiosis were identified. Then, a total of 29,752 differentially expressed genes (DEGs) were identified between the ovary and testis tissue, including 18,570 up-regulated genes and 11,182 down-regulated genes in the testis compared with the ovary in the discus fish. Among these DEGs, some of the male-enhanced genes included the following the genes. *Dmrt1* (double-sex and mab-3-related transcription factor, which has been demonstrated to be the duplicated homologs of the medaka *(Oryzias latipes)*. The *DMY* gene [35], which was first found as sex-determination gene in non-mammalian vertebrates, and has a conservative role in sex determination and differentiation of vertebrates as an ancestral function. *Dmrt2*, which is more likely to be involved in male gonadal development or maintenance of gonadal function in *Chlamys nobilis* [36]. *Cyp17a1*, which is the qualitative regulator of steroidogenesis [37], whose expression levels in the testis was higher compared to the ovary, which indicates that *Cyp17a1* may be involved in testicular formation during sex differentiation [38]. *Amh*, a male-specific gene, is derived from the Sertoli cells at the initiation of testis differentiation, participates in steroidogenic pathway to produce testosterone, or negatively controls oestrogen production [39]. Their expression levels were significantly higher in the testis than in the ovary using the RNA-Seq and qRT-PCR methods, which indicates that they may play key roles in regulating testis development in the discus fish.

Additionally, some of the female-enhanced genes were identified to be various ovary marker genes. These include the following several genes. *Sox3*, which is highly conserved in fish sex differentiation pathways [40], plays a key role in regulating gametogenesis and gonad differentiation of vertebrates, and previous studies have shown that it might have more important effect in oogenesis than in spermatogenesis [41]. *Cyp19a1*, a Cytochrome P450 aromatase, that can catalyse the conversion of androgens to oestrogens, and plays dual roles in regulating testicular development during the initial period of sexual differentiation and later in ovarian development during the natural sex change in the protandrous black porgy *(Acanthopagrus schlegeli)* [42]; In pejerrey *(Odontesthes bonariensis)*, the tissue distribution analysis of *cyp19a1* mRNA in adult fish revealed high expression in the ovary, which is involved in the process of ovarian formation [43]. *Dnd*, as an ovarian marker, plays an essential role during female gametogenesis and embryo development in pigs [44]. However, *Sox9* expressed no difference between the testis and ovary in the discus fish. A possible reason is that *Sox9* participates in an early stage of gonadal development and is identified as an early signal of ovarian differentiation [45].

We performed a comprehensive annotation and comparative analysis of miRNA using high-throughput sequencing and bioinformatics methods, and identified 47 differentially expressed miRNAs between the testis and the ovary in the discus fish, including 31 significantly up-regulated miRNAs and 16 significantly down-regulated miRNAs with testis compared to the ovary. In this study, the results from miRNA RNA-Seq sequencing indicated that miR-7641, miR-205a, miR-181a-5p, miR-143-3p, miR-145-3p and miR-129-5p were differentially expressed between testis and ovary in the discus fish. miRNAs play crucial roles in a variety of biological processes via regulating expression of their target genes at the mRNA level. There is increasing evidence that miRNAs play an important role in regulating biological process, while miRNAs targeting mRNA is a key part of understanding their role in gene regulation networks [46, 47]. In this study, 41 miRNA-mRNA interaction pairs were identified with 15 DEGs mRNA targeted by 24 differentially expressed miRNAs.

miRNAs play crucial roles in a variety of biological processes via regulating expression of their target genes at the mRNA level. The miRNA-mRNA interaction pair with miRNA targeting mRNA is a key part of understanding miRNA function. In this study, 41 miRNA-mRNA interaction pairs were identified with 15 differentially expressed genes targeted by 24 differentially expressed miRNAs using miRNA target prediction method. In these pairs, miR-2187-3p, miR-24-3p, miR-181a, miR-181-5p, miR-101a, miR-218-5p, miR-217 and miR-7133 can target *Hsd11β2*, which is involved in the steroid hormone biosynthesis pathway, indicating that these miRNAs might play an important role in regulating steroid hormone biosynthesis; miR-181a can also target *Hsd17β10*, which is involved in the steroid hormone biosynthesis pathway, and both were down-regulated in testes, indicating that miR-181a might exert different functions on regulating steroid hormone biosynthesis between ovaries and testes. Otherwise, previous studies showed that *Dmrt1* was predominantly expressed in spermatogonia, spermatocytes and spermatids, as well as in Sertoli cells, indicating that *Dmrt1* plays an important role in spermatogenesis in *Halobatrachus didactylus* [48]. In the discus fish, *Dmrt1* was mainly expressed in the testes, while weakly in the ovaries, and the *Dmrt1* gene was targeted by miR-129-5p, miR-27b-3p, miR-27e, miR-152, miR-27b-3p, miR-17-3p and miR-460 in our miRNA target prediction data, which indicates that these might be involved in regulating spermatogenesis and testis development of the discus fish. Additionally, miR-129-5p was highly expressed in the ovary, weakly expressed in the testis, and could target *Dmrt1*, GHR (growth hormone receptor) and RERG (ras-related and oestrogen-regulated growth inhibitor), indicating that miR-129-5p might play important roles in regulating growth development and physiological function of the ovary in the discus fish. Previous studies have shown that *zp3* is found on the extracellular matrix of oocytes and acts as a receptor for mammalian sperm binding. In this study, we found that *zp3* can be targeted by miR-7641 and miR-205a in the miRNAs target prediction data, and miR-7641 and miR-205a were lowly expressed in the ovaries of the discus fish, while highly in the testes, which indicating that lower expression of the two miRNAs in the ovaries might be necessary for expression of the *zp3* gene, playing an important role in regulating the biological process of the ovary binding sperm. Therefore, in the discus fish, miRNAs can play important roles in regulating steroid hormone synthesis, gonadal physiological function and sex development by regulating their target genes. However, the real function of these miRNAs needs to be confirmed in the gonad of the discus fish in the further studies.

In summary, the discus fish *(S. aequifasciatus)* is one of the most beautiful ornamental fish species with brilliant colours and pretty disc-shaped body. However, there are hardly any differences between male and female fish. In our study, we found some differentially expressed genes and miRNAs between the ovary and testis, and predicted target relations between genes and miRNAs using miRNA target prediction methods, and these differentially expressed genes and miRNAs were involved in regulating gonadal development, gametogenesis, physiological function of ovary and testis. These results can help us further understand the mechanism of gonad development between female and male discus fish and provide data for further study of the sex development of the discus fish in the future.

## Acknowledgements

This study presented in the manuscript was funded by the Key Project of Developing Agriculture through Science and Technology of Shanghai Municipal Agricultural Commission (2015–19), China; the China Postdoctoral Science Foundation (2017M621433); and the Doctoral Scientific Research Foundation of Shanghai Ocean University (A2-0203-17-100306). LC-Bio (Hangzhou, China) carried out the Illumina RNA-Seq sequencing of miRNAs and mRNA libraries of the testes and ovaries in the discus fish.

## Author contributions statement

Z.Z.C. and Y.S.F. designed the study. Y.S.F. and Z.X. performed the bioinformatics analysis and verification experiments, and wrote the manuscript. Z.Z.C. and J.Z.G. supervised the progress of the project. B.W. provide important suggestions about the manuscript writing and collect the samples. All authors reviewed the manuscript.

## Additional information

**Supplementary information** includes Supplementary Figures and Supplementary Tables accompanying this paper.

## Supporting information

### Supplementary Figures

**Supplementary Figure 1.** Functional classification of assembled unique sequences based on gene ontology (GO) terms: molecular function, cellular component, and biological process.

**Supplementary Figure 2.** Functional categories of KOG database in discus fish.

**Supplementary Figure 3.** KEGG Pathway Classification of the gonad in discus fish.

### Supplementary Tables

**Supplementary Table S1.** miRNA and mRNA Primers used in this study for qRT-PCR validation. (XLSX)

**Supplementary Table S2.** The differential expression mRNA between testis and ovary. (XLSX)

**Supplementary Table S3.** GO functional annotation of DEGs for testis and ovary. (XLSX)

**Supplementary Table S4.** KEGG enrichment annotation of DEGs for testis and ovary. (XLSX)

**Supplementary Table S5.** The differential expression miRNA between testis and ovary. (XLSX)

**Supplementary Table S6.** The data for qRT-PCR and RNA-Seq of mRNAs and miRNA. (XLSX)

**Supplementary Table S7.** 41 miRNA-mRNA interaction pairs involved in sex development and maintenance between testis and ovary comparisons respectively. (XLSX)

## References

1. Webster KA, Schach U, Ordaz A, Steinfeld JS, Draper BW, Siegfried KR. *Dmrt1* is necessary for male sexual development in zebrafish. Dev Biol. 2017;422(1):33–46. doi: 10.1016/j.ydbio.2016.12.008. PubMed PMID: 27940159.

2. Baroiller JF, Guiguen Y. Endocrine and environmental aspects of sex differentiation in gonochoristic fish. Exs. 2001;(91):177–201. PubMed PMID: 11301598.

3. Nagahama Y. Molecular mechanisms of sex determination and gonadal sex differentiation in fish. Fish Physiol Biochem. 2005;31(2–3):105–9. doi: 10.1007/s10695-006-7590-2. PubMed PMID: 20035442.

4. Baroiller JF, D’Cotta H, Saillant E. Environmental effects on fish sex determination and differentiation. Sexual development: genetics, molecular biology, evolution, endocrinology, embryology, and pathology of sex determination and differentiation. 2009;3(2–3):118–35. doi: 10.1159/000223077. PubMed PMID: 19684457.

5. Shen ZG, Wang HP. Molecular players involved in temperature-dependent sex determination and sex differentiation in Teleost fish. Genetics, selection, evolution : GSE. 2014;46:26. doi: 10.1186/1297-9686-46-26. PubMed PMID: 24735220; PubMed Central PMCID: PMC4108122.

6. Sinclair AH, Berta P, Palmer MS, Hawkins JR, Griffiths BL, Smith MJ, et al. A gene from the human sex-determining region encodes a protein with homology to a conserved DNA-binding motif. Nature. 1990;346(6281):240–4. doi: 10.1038/346240a0. PubMed PMID: 1695712.

7. Matsuda M, Nagahama Y, Shinomiya A, Sato T, Matsuda C, Kobayashi T, et al. DMY is a Y-specific DM-domain gene required for male development in the medaka fish. Nature. 2002;417(6888):559–63. doi: 10.1038/nature751. PubMed PMID: 12037570.

8. Nanda I, Kondo M, Hornung U, Asakawa S, Winkler C, Shimizu A, et al. A duplicated copy of DMRT1 in the sex-determining region of the Y chromosome of the medaka, Oryzias latipes. Proc Natl Acad Sci U S A. 2002;99(18):11778–83. doi: 10.1073/pnas.182314699. PubMed PMID: 12193652; PubMed Central PMCID: PMC129345.

9. Huang S, Ye L, Chen H. Sex determination and maintenance: the role of DMRT1 and FOXL2. Asian journal of andrology. 2017;19(6):619–24. doi: 10.4103/1008-682X.194420. PubMed PMID: 28091399.

10. de Rooij DG, Russell LD. All you wanted to know about spermatogonia but were afraid to ask. Journal of Andrology. 2000;21(6):776–98.

11. Shiraishi K. HSF Is Required for Gametogenesis: Springer Japan; 2016.

12. Cabas I, Chaves-Pozo E, García-Alcázar A, Meseguer J, Mulero V, García-Ayala A. The effect of 17α-ethynylestradiol on steroidogenesis and gonadal cytokine gene expression is related to the reproductive stage in marine hermaphrodite fish. Marine Drugs. 2013;11(12):4973–92.

13. Senthilkumaran B, Sudhakumari CC, Wang DS, Sreenivasulu G, Kobayashi T, Kobayashi HK, et al. Novel 3β-hydroxysteroid dehydrogenases from gonads of the Nile tilapia: phylogenetic significance and expression during reproductive cycle. Molecular & Cellular Endocrinology. 2009;299(2):146–52.

14. Ambros V. The functions of animal microRNAs. Nature. 2004;431(7006):350–5. doi: Doi 10.1038/Nature02871. PubMed PMID: WOS:000223864000051.

15. Bartel DP. MicroRNAs: Target Recognition and Regulatory Functions. Cell. 2009;136(2):215.

16. Mardis ER. The impact of next-generation sequencing technology on genetics. Trends Genet. 2008;24(3):133–41. Epub 2008/02/12. doi: 10.1016/j.tig.2007.12.007. PubMed PMID: 18262675.

17. Bartel DP. MicroRNAs: genomics, biogenesis, mechanism, and function. Cell. 2004;116(2):281–97. Epub 2004/01/28. PubMed PMID: 14744438.

18. Niwa R, Slack FJ. The evolution of animal microRNA function. Current opinion in genetics & development. 2007;17(2):145–50. doi: 10.1016/j.gde.2007.02.004. PubMed PMID: 17317150.

19. Sempere LF, Cole CN, McPeek MA, Peterson KJ. The phylogenetic distribution of metazoan microRNAs: insights into evolutionary complexity and constraint. Journal of experimental zoology Part B, Molecular and developmental evolution. 2006;306(6):575– 88. doi: 10.1002/jez.b.21118. PubMed PMID: 16838302.

20. Heimberg AM, Sempere LF, Moy VN, Donoghue PC, Peterson KJ. MicroRNAs and the advent of vertebrate morphological complexity. Proc Natl Acad Sci U S A. 2008;105(8):2946–50. doi: 10.1073/pnas.0712259105. PubMed PMID: 18287013; PubMed Central PMCID: PMC2268565.

21. Tao W, Sun L, Shi H, Cheng Y, Jiang D, Fu B, et al. Integrated analysis of miRNA and mRNA expression profiles in tilapia gonads at an early stage of sex differentiation. Bmc Genomics. 2016;17(1):328.

22. Juanchich A, Cam AL, Montfort J, Guiguen Y, Bobe J. Identification of Differentially Expressed miRNAs and Their Potential Targets During Fish Ovarian Development1. Biology of Reproduction. 2013;88(5):128.

23. Patel RK, Jain M. NGS QC Toolkit: a toolkit for quality control of next generation sequencing data. Plos One. 2012;7(2):e30619. Epub 2012/02/09. doi: 10.1371/journal.pone.0030619. PubMed PMID: 22312429; PubMed Central PMCID: PMCPMC3270013.

24. Grabherr MG, Haas BJ, Yassour M, Levin JZ, Thompson DA, Amit I, et al. Full-length transcriptome assembly from RNA-Seq data without a reference genome. Nat Biotechnol. 2011;29(7):644–U130. doi: 10.1038/nbt.1883. PubMed PMID: WOS:000292595200023.

25. Ashburner M, Ball CA, Blake JA, Botstein D, Butler H, Cherry JM, et al. Gene Ontology: tool for the unification of biology. Nat Genet. 2000;25(1):25–9. doi: Doi 10.1038/75556. PubMed PMID: WOS:000086884000011.

26. Tatusov RL, Fedorova ND, Jackson JD, Jacobs AR, Kiryutin B, Koonin EV, et al. The COG database: an updated version includes eukaryotes. Bmc Bioinformatics. 2003;4:41. Epub 2003/09/13. doi: 10.1186/1471-2105-4-41. PubMed PMID: 12969510; PubMed Central PMCID: PMCPMC222959.

27. Kanehisa M, Araki M, Goto S, Hattori M, Hirakawa M, Itoh M, et al. KEGG for linking genomes to life and the environment. Nucleic Acids Res. 2008;36:D480–D4. doi: 10.1093/nar/gkm882. PubMed PMID: WOS:000252545400086.

28. Kozomara A, Griffiths-Jones S. miRBase: annotating high confidence microRNAs using deep sequencing data. Nucleic Acids Res. 2014;42(Database issue):D68–73. Epub 2013/11/28. doi: 10.1093/nar/gkt1181. PubMed PMID: 24275495; PubMed Central PMCID: PMCPMC3965103.

29. Fu Y, Shi Z, Wu M, Zhang J, Jia L, Chen X. Identification and differential expression of microRNAs during metamorphosis of the Japanese flounder (Paralichthys olivaceus). Plos One. 2011;6(7):e22957. Epub 2011/08/06. doi: 10.1371/journal.pone.0022957. PubMed PMID: 21818405; PubMed Central PMCID: PMCPMC3144956.

30. Shi Y, Yang F, Wei S, Xu G. Identification of Key Genes Affecting Results of Hyperthermia in Osteosarcoma Based on Integrative ChIP-Seq/TargetScan Analysis. Med Sci Monit. 2017;23:2042–8. Epub 2017/04/30. PubMed PMID: 28453502; PubMed Central PMCID: PMCPMC5419091.

31. John B, Enright AJ, Aravin A, Tuschl T, Sander C, Marks DS. Human MicroRNA targets. PLoS Biol. 2004;2(11):e363. Epub 2004/10/27. doi: 10.1371/journal.pbio.0020363. PubMed PMID: 15502875; PubMed Central PMCID: PMCPMC521178.

32. Li B, Ruotti V, Stewart RM, Thomson JA, Dewey CN. RNA-Seq gene expression estimation with read mapping uncertainty. Bioinformatics. 2010;26(4):493–500. Epub 2009/12/22. doi: 10.1093/bioinformatics/btp692. PubMed PMID: 20022975; PubMed Central PMCID: PMCPMC2820677.

33. Li B, Dewey CN. RSEM: accurate transcript quantification from RNA-Seq data with or without a reference genome. Bmc Bioinformatics. 2011;12:323. Epub 2011/08/06. doi: 10.1186/1471-2105-12-323. PubMed PMID: 21816040; PubMed Central PMCID: PMCPMC3163565.

34. Cer RZ, Herreragaleano JE, Anderson JJ, Bishoplilly KA, Mokashi VP. miRNA Temporal Analyzer (mirnaTA): a bioinformatics tool for identifying differentially expressed microRNAs in temporal studies using normal quantile transformation. GigaScience,3,1(2014-10-13). 2014;3(1):20.

35. Matsuda M. Sex determination in fish: Lessons from the sex-determining gene of the teleost medaka, Oryzias latipes. Development Growth & Differentiation. 2003;45(5-6):397–403.

36. Shi Y, Wang Q, He M. Molecular identification of dmrt2 and dmrt5 and effect of sex steroids on their expressions in Chlamys nobilis. Aquaculture. 2014;s 426–427(1):21–30.

37. Missaghian E, Kempná P, Dick B, Hirsch A, Alikhanikoupaei R, Jégou B, et al. Role of DNA methylation in the tissue-specific expression of the CYP17A1 gene for steroidogenesis in rodents. Journal of Endocrinology. 2009;202(1):99–109.

38. Sakurai N, Maruo K, Haraguchi S, Uno Y, Oshima Y, Tsutsui K, et al. Immunohistochemical detection and biological activities of CYP17 (P450c17) in the indifferent gonad of the frog Rana rugosa. Journal of Steroid Biochemistry & Molecular Biology. 2008;112(1–3):5–12.

39. Poonlaphdecha S, Pepey E, Huang SH, Canonne M, Soler L, Mortaji S, et al. Elevated amh gene expression in the brain of male tilapia (Oreochromis niloticus) during testis differentiation. Sexual Development Genetics Molecular Biology Evolution Endocrinology Embryology & Pathology of Sex Determination & Differentiation. 2011;5(1):33–47.

40. Takehana Y, Matsuda M, Myosho T, Suster ML, Kawakami K, Shin-I T, et al. Co-option of Sox3 as the male-determining factor on the Y chromosome in the fish Oryzias dancena. Nature Communications. 2014;5:4157.

41. Bo Y, Li Z, Yang W, Wei X, Gui JF. Differential expression and dynamic changes of SOX3 during gametogenesis and sex reversal in protogynous hermaphroditic fish. Journal of Experimental Zoology Part A Ecological Genetics & Physiology. 2007;307A(4):207–19.

42. Wu GC, Tomy S, Nakamura M, Chang CF. Dual roles of cyp19a1a in gonadal sex differentiation and development in the protandrous black porgy, Acanthopagrus schlegeli. Biology of Reproduction. 2008;79(6):1111.

43. Karube M, Fernandino JI, Stroblmazzulla P, Strüssmann CA, Yoshizaki G, Somoza GM, et al. Characterization and expression profile of the ovarian cytochrome P-450 aromatase (cyp19A1) gene during thermolabile sex determination in pejerrey, Odontesthes bonariensis. Journal of Experimental Zoology Part A Ecological Genetics & Physiology. 2007;307A(11):625–36.

44. Yang CX, Wright EC, Ross JW. Expression of RNA-binding proteins DND1 and FXR1 in the porcine ovary, and during oocyte maturation and early embryo development. Molecular Reproduction & Development. 2012;79(8):541–52.

45. Berbejillo J, Martinez-Bengochea A, Bedó G, Vizziano-Cantonnet D. Expression of dmrt1 and sox9 during gonadal development in the Siberian sturgeon (Acipenser baerii). Fish Physiology & Biochemistry. 2013;39(1):91–4.

46. Jing J, Wu J, Liu W, Xiong S, Ma W, Zhang J, et al. Sex-Biased miRNAs in Gonad and Their Potential Roles for Testis Development in Yellow Catfish. Plos One. 2014;9(9):e107946.

47. Zhang X, Yuan L, Li L, Jiang H, Chen J. Conservation, sex-biased expression and functional annotation of microRNAs in the gonad of Amur sturgeon (Acipenser schrenckii). Comparative Biochemistry & Physiology Part D Genomics & Proteomics. 2016;18:54.

48. Ubeda-Manzanaro M, Merlo MA, Ortiz-Delgado JB, Rebordinos L, Sarasquete C. Expression profiling of the sex-related gene Dmrt1 in adults of the Lusitanian toadfish Halobatrachus didactylus (Bloch and Schneider, 1801). Gene. 2014;535(2):255.

